# Polyunsaturated fatty acids and their endocannabinoid-related metabolites activity at human TRPV1 and TRPA1 ion channels expressed in HEK-293 cells

**DOI:** 10.1101/2024.09.25.615078

**Authors:** Atnaf A Abate, Marina J Santiago, Alf Garcia-Bennett, Mark Connor

## Abstract

**Background:** Polyunsaturated fatty acids (PUFAs), particularly Omega-3 (ω-3) and Omega-6 (ω-6) PUFAs, may exert neuroprotective effects via the endocannabinoid system (ECS) and are promoted as brain health supplements. However, despite their potential role in endocannabinoid biosynthesis, the impact of PUFAs on ion channels such as TRPV1 and TRPA1, which are modulated by endocannabinoids, remains incompletely understood. Furthermore, the potential *in vitro* actions of ω-6 to ω-3 PUFA combined in the ratios available in supplements remains uncertain. Therefore, the objective of this study is to evaluate the functional activity of individual PUFAs, their combination in a specific ratio, and their endocannabinoid-related derivatives on TRPV1 and TRPA1 ion channels.

**Methodology:** We employed a fluorescent Ca-sensitive dye in HEK-293 Flp-In T-REx cells expressing human TRPV1, TRPA1, or an empty vector to measure changes in intracellular calcium concentration ([Ca^2+]^_i_).

**Results:** Capsaicin and certain PUFA derivatives such as DHEA, γ-LEA and AEA stimulate TRPV1 activity directly, whereas EPA, DHA, γ-LA, and their 9:3:1 ratio triggered TRPV1 response via a mechanism dependent on prior exposure to phorbol ester. Similarly, cinnamaldehyde and selected PUFA derivatives such as EPEA, DHEA, γ-LEA, 2-AG, 2-AG ether and AEA triggered TRPA1 response, with EPA, DHA, γ-LA, and the 9:3:1 ratio showing significant effects at higher concentrations.

**Conclusions:** PUFAs alone and their combined form in 9:3:1 ratio stimulate TRPA1 activity, whereas their metabolites trigger both TRPV1 and TRPA1 response. These findings suggest new avenues to explore for research into potential mechanisms underlying the neurological benefits of PUFAs and their metabolites.

## INTRODUCTION

Polyunsaturated fatty acids (PUFAs) are crucial for maintaining brain function and has been used as supplements with claims of improving brain health (Bentsen 2017; Bourre 2009). In preclinical models, omega-3 (ω-3) and omega-6 (ω-6) PUFAs have demonstrated neuroprotective effects via the endocannabinoid system (ECS) (Dyall 2017; Freitas et al. 2018).

The ECS is an essential part of the central nervous system (CNS) (Skaper & Di Marzo 2012). Its core components include the lipid derivatives of PUFA’s, endocannabinoids (eCB), enzymes regulating eCB synthesis and breakdown, and cannabinoid receptors CB1 and CB2 (DeMesa et al. 2021; Fezza et al. 2014; Lu & Mackie 2016). Additionally, other G protein-coupled receptors like GPR55 (Lauckner et al. 2008; Yang et al. 2016), GPR3, GPR119 (Davis 2022) and GPR120 (Im 2009) are the potential member of ECS. Peroxisome-proliferator activated receptors (PPARs) and transient receptor potential (TRP) ion channels, are also activated by various cannabinoid ligands, including eCBs (DeMesa et al. 2021; Lu & Mackie 2016).

TRP channels are membrane proteins involved in sensing and responding to chemical and physical stimuli. They are integral to neural signalling processes related to various sensory perceptions including nociception (Julius 2013; Muller et al. 2019; Sawamura et al. 2017; Zhang et al. 2023). Specific channels within the TRP family, such as TRPV1, TRPV2, TRPV3, TRPV4, TRPA1, and TRPM8, have been identified as responsive to endogenous, phyto-, and synthetic cannabinoids (Muller et al. 2019). It is also reported that, these channels may contribute to eCB signalling, especially within the brain (Nilius & Owsianik 2011). However, the potential modulation of these ion channels by ω-3 and ω-6 fatty acids derived eCBs has not been fully defined (Petermann et al. 2022).

Dietary PUFAs activate TRP channels, with eicosapentaenoic acid (EPA) and docosahexaenoic acid (DHA) shown to activate TRPV1(Ciardo & Ferrer-Montiel 2017; Matta et al. 2007) and TRPA1 (Ciardo & Ferrer-Montiel 2017; Motter & Ahern 2012; Redmond et al. 2014) ion channels. These PUFAs also serve as major precursors for eCB biosynthesis (Komarnytsky et al. 2021), with resulting eCBs activating TRPV1 and TRPA1 ion channels. It has been reported that 2-arachidonoylglycerol (2-AG), anandamide (AEA) (Zygmunt et al. 2013; Zygmunt et al. 1999), and N-arachidonoyldopamine (NADA) (Huang et al. 2002; Raboune et al. 2014) activate TRPV1, while AEA activates TRPA1 (Redmond et al. 2014). However, there are also other ethanolamides or glycerol conjugates of PUFAs that may function as eCBs or ligands for related receptors; but their roles and targets in the brain remain unclear (Bosch-Bouju & Laye 2016; Witkamp 2016). The impact of the ω-6 fatty acid such as γ-linolenic acid (γ-LA) and its ethanolamine derivative on TRPV1 and TRPA1 ion channels has also not been examined.

Dietary intake affects brain PUFA levels (Dyall 2017), and the ω-6 to ω-3 ratio in the current Western diet (approximately 20:1) is linked to various health conditions including autoimmune and inflammatory diseases (Patel et al. 2022; Simopoulos 2002). One possible reason could be that ω-3 long-chain fatty acids such as EPA, DHA as well as some ω-6 derived fatty acids, e.g., γ-LA particularly dihomo - γ-linolenic acid, are important for production of anti-inflammatory eicosanoids while ω-6 derived fatty acids, mainly AA, are crucial to produce pro-inflammatory eicosanoids. The presence of higher amounts of pro-inflammatory eicosanoids such as AA also interferes the synthesis of anti-inflammatory eicosanoids by competing at the active site of the enzyme, cyclooxygenase (COX) (Calder 2010). Therefore, to mitigate risks linked with excessive ω-6 PUFA consumption, maintaining a balanced ω-6 to ω-3 ratio of 1:1 to 5:1 is suggested (Patel et al. 2022). Studies have also indicated that the ratio of ω-6 to ω-3 fatty acids in tissues is more important for health benefits than their absolute levels (lshweki et al. 2015) and combining ω-6 and ω-3 PUFAs has shown positive health effects, especially in children with Developmental coordination disorder (DCD) (Richardson & Montgomery 2005). Moreover, it is noted that maintaining an equilibrium between ω-6 and ω-3 PUFAs in a healthy diet yields favourable effects on inflammation and other physiological mechanisms (Gomez-Candela et al. 2011). Nevertheless, common agreement regarding the ideal ratio remains elusive. Determining the optimum ratio of ω-3 PUFAs, EPA, and DHA for desired health benefits also remains uncertain (Djuricic & Calder 2021; Gomez-Candela et al. 2011; Mukhametove et al. 2022).

The objective of this study is to evaluate the individual PUFAs (EPA, DHA, and γ-LA), their combined form in a 9:3:1 ratio, and their endocannabinoid-related derivatives on the functional activity of human TRPV1 and TRPA1 ion channels, to develop a more complete picture of the mode of actions of these important dietary molecules.

## METHODOLOGY

### Cell culture

Flp-In T-REx HEK-293 cells (Life Technologies, Mulgrave, Victoria, Australia), stably transfected with human TRPV1 (hTRPV1), TRPA1 (hTRPA1) cDNA (GenScript, Piscataway, NJ) (Heblinski et al. 2020) or an empty pcDNA5/FR/TO vector, were maintained in Dulbecco’s Modified Eagle’s Medium (DMEM) with 10% fetal bovine serum (FBS) (Gibco, # 10099-141), 100 U penicillin and 100 μg streptomycin ml^−1^ (1% P/S) (Gibco, # 15140-122, Life Technologies, USA), 80 μg.ml^−1^ hygromycin B (Invivogen, cat # ant-hg-1), and 15μg.ml^−1^ blasticidin (Invivogen; San Diego, CA, USA, cat # ant-bl-1). The cells were incubated in a 5% CO_2_ humidified atmosphere at 37°C. Upon reaching approximately 90% confluence in 75cm^2^ flasks, they were trypsinized and transferred to poly-D-lysine coated 96-well plates (Corning, Castle Hill, NSW, Australia) in L-15 medium supplemented with 1% FBS, 100 U penicillin and 100 μg streptomycin ml^−1^ (1% P/S), and 15 mM glucose (80 μL volume per well). After an overnight incubation (maximum of 16 hours) in humidified room air at 37°C, TRPV1 and TRPA1 receptor expression was induced 4 hours before experimentation by adding to each well 10 μL tetracycline solution to a final concentration of 1 μg.ml^−1^.

### Calcium Assay

Intracellular calcium [Ca]_i_ levels were assessed using the calcium 5 kit from Molecular Devices (# R8186, Sunnyvale, CA, USA) on a FlexStation 3 Microplate Reader (Molecular Devices, Sunnyvale, CA, USA). Calcium 5 dye was dissolved in Hank’s Balanced Salt Solution (HBSS) with the composition of (in mM): NaCl 145, CaCl_2_ 1.26, MgCl_2_ 0.493, HEPES 22, Na_2_HPO_4_ 0.338, NaHCO_3_ 4.17, KH_2_PO_4_ 0.441, MgSO_4_ 0.407 and glucose 1mg/ml (pH adjusted to 7.4, osmolarity = 315 ± 15 mosmol) and used at 50% of the manufacturer’s suggested concentration. Probenecid (Biotium, cat # 50027) which helps to prevent expulsion of calcium indicator from the cells was added to a final concentration of 1.25 mM. 90μL of the dye were loaded into each well of the plate and incubated for 1 hour before reading in the FlexStation 3 at 37°C. Fluorescence was recorded every 2 seconds (λ excitation = 485 nm, λ emission = 525 nm) for 5 minutes. After 1 minute of baseline recording, 20 μL of the drug, dissolved in HBSS with 1% Dimethyl Sulfoxide (DMSO) (Sigma-Aldrich, Germany, cat # D2650) was added (final DMSO concentration was 0.1% in well).

### Drugs and reagents

All drugs were prepared in DMSO at a concentration of 30 mM and stored at -30°C/-80°C. Freshly thawed aliquots were used in each experiment and diluted in HBSS containing 0.1% bovine serum albumin (BSA) (Sigma-Aldrich, US, # A7030). Due to limitations in the solubility of fatty acids and their derivatives, the highest concentration tested was 30 μM. EPA, DHA, γ-LA, and their endocannabinoid related derivatives and glycerol conjugates were procured from Cayman Chemical (Ann Arbor, MI, USA) while cinnamaldehyde (CA) was obtained from Merck (Castle Hill, NSW, Australia). The 9:3:1 ratio were prepared by combining 9 % of EPA, 3 % of DHA and 1 % of γ-LA dissolved in DMSO, so that, the highest concentration tested was 30:10:3 µM. In this study, we explored the effects of a 9:3:1 ratio of EPA, DHA, and γ-LA, which reflects the composition found in commercially available omega-3 supplements. This ratio was chosen because an excess of EPA compared to GLA is prevalent in these supplements, designed to maximize the anti-inflammatory benefits associated with omega-3 fatty acids (Baker et al. 2016). All reagents for tissue culture were sourced from Merck or Life Technologies (Mulgrave, Victoria, Australia). Capsaicin (Caps) was obtained from Tocris Bioscience, Bristol (# 404-86-4) and capsazepine was from Merck, USA (# 138977-28-3). PAR-1 agonist peptide (Thr-Phe-Leu-Leu-Arg-NH_2_, # 2660) was obtained from Auspep (Tullamarine, Victoria, Australia).

### Data analysis

The response to agonists was expressed as a percentage change from the average baseline measurement (1 minute before adding drug). Changes in fluorescence resulting from the addition of solvent were subtracted before normalization to baseline. Concentration-response curves, E_max_, EC_50_ values and Hill slope were determined using a three-parameter logistic equation (GraphPad Prism, San Diego, CA). Results are presented as the mean ± standard error of the mean (SEM) from at least six separate experiments conducted in duplicate, unless otherwise stated. Concentration-response curves (CRC) of the positive controls, capsaicin and cinnamaldehyde, were performed each day for comparative analysis. Where appropriate, unpaired Student’s t-test were used to compare the responses of individual compounds in different conditions, while a One-way ANOVA followed by Dunnett’s multiple comparisons test was used to assess potential differences in responses elicited by a range of compounds. P < 0.05 was considered statistically significant.

## RESULTS

Application of docosahexaenoyl ethanolamide (DHEA), γ-LEA, AEA, 2-linoleoyl glycerol (2-LG), NADA, and capsaicin at 10 µM produced an elevation [Ca]_i_ in HEK-293 TRPV1 cells that was significantly greater than that produced in HEK-293 EV cells (Figure 1, Table 1). However, EPA, DHA, γ-LA, their 9:3:1 ratio, as well as EPEA, 2-AG and 2-AG ether did not produce a notable change in [Ca]_i_ in either cell line, and there was no difference between the responses in HEK-293 TRPV1 cells and those seen in HEK-293 EV cells (Table 1).

**Figure 1.**
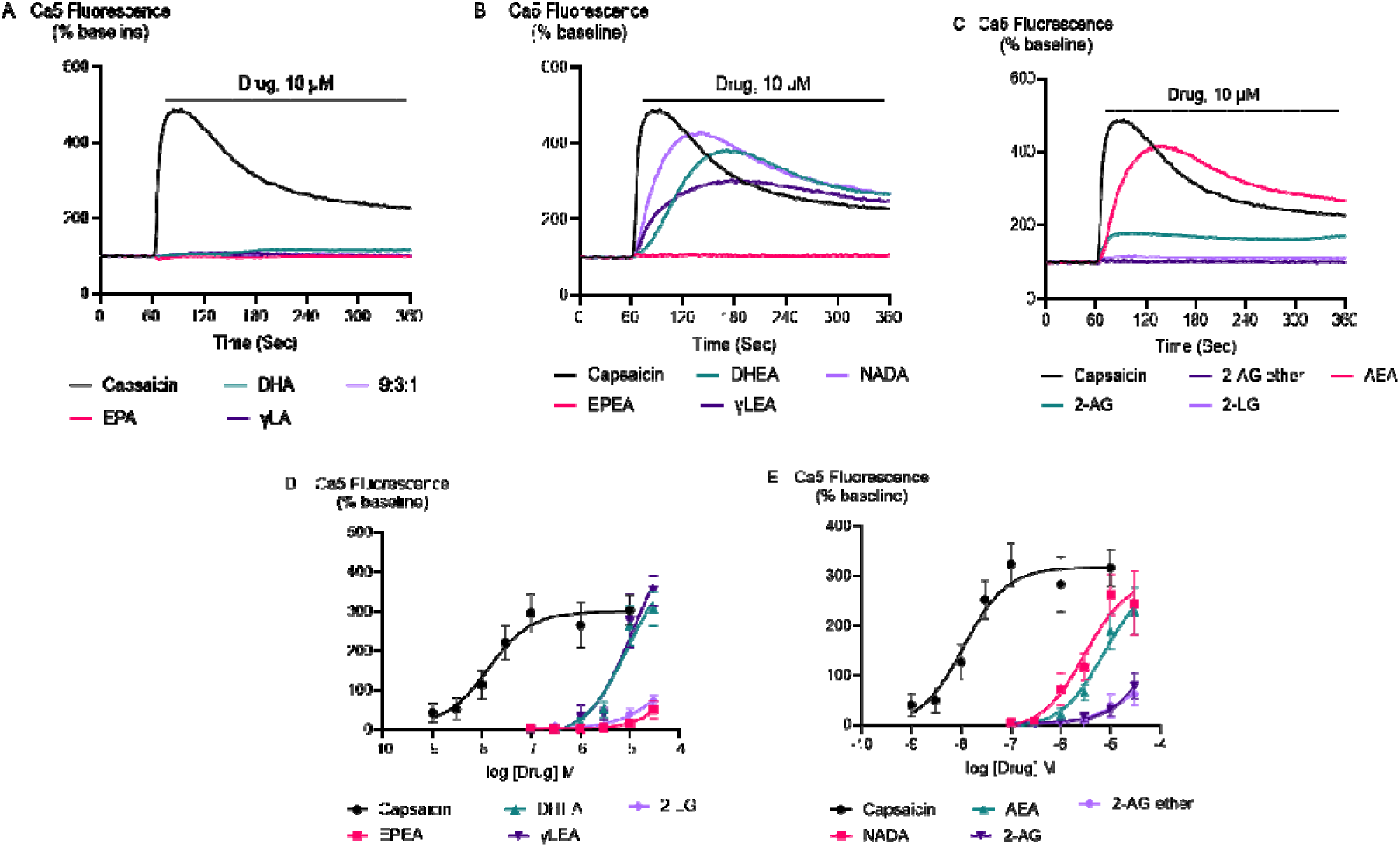
Traces of PUFAs and their endocannabinoid related metabolites at 10 µM (A, B and C) and concentration-response curves (CRC) for endocannabinoid-related metabolites and capsaicin (D and E) in HEK-293 TRPV1 expressing cells. In A), B) and C) drugs were added for the duration of the bar. The CRCs were fitted using a three-parameter logistic equation, each data point represents the mean ± SEM from at least 6 independent experiments conducted in duplicate.

**Table 1.** Change of Ca5 fluorescence induced by PUFAs and their endocannabinoid related metabolites in HEK-293 EV, TRPV1 and TRPA1 expressing cells.

| DRUG | Mean± SEM | | |
| --- | --- | --- | --- |
|  | EV | TRPV1 | TRPA1 |
| EPA□ | 14 ± 5□ | 16 ± 5□ | 58 ± 42□ |
| DHA□ | 11 ± 2□ | 11 ± 2□ | 129 ± 81□ |
| $\gamma$ - LA□ | 9 ± 2□ | 10 ± 1□ | 112 ± 61□ |
| 9:3:1□ | 9 ± 1 | 11 ± 2□ | 85 ± 45□ |
| EPEA□ | 3 ± 1□ | 15 ± 8□ | 50 ± 40□ |
| DHEA□ | 10 ± 3 | 266 ± 51*□ | 107 ± 50*□ |
| γ-LEA□ | 11 ± 2 | 277 ± 67*□ | 87 ± 32*□ |
| AEA□ | 10 ± 2 | 188 ± 36*□ | 78 ± 40□ |
| 2-AG□ | 4 ± 2□ | 30 ± 12□ | 6 ± 1□ |
| 2-AG ether□ | 8 ± 2□ | 39 ± 24□ | 91 ± 40□ |
| 2-LG□ | 4 ± 1 | 41 ± 13*□ | 6 ± 2□ |
| NADA□ | 6 ± 2 | 262 ± 40 *□ | -□ |
| Par-1 (100 μM)□ | 463 ± 33 | -□ | -□ |
| Capsaicin□□ | 12 ± 2 | □350 ± 33 *□ | -□ |
| Cinnamaldehyde (300 μM)□ | 9 ± 6 | -□ | □438 ± 29 *□ |

Drugs were applied at 10 μM. Responses are expressed as a % change in fluorescence from baseline.
All experiments represent mean ± SEM of at least 6 independent experiments, except for Cinnamaldehyde in EV cells, which was n=4. An unpaired t-test was conducted to compare the response of PUFAs and their endocannabinoid-related metabolites in HEK-293 EV cell with that of HEK-293 TRPV1 and TRPA1 expressing cells. “\*” Indicates P < 0.05 compared to empty vector cells.

The effects of the positive control and endocannabinoid-related metabolites of PUFA’s in HEK-293 TRPV1 expressing cells were concentration-dependent. Capsaicin, a prototypic TRPV1 agonist, increased Ca5-dye fluorescence, peaking at a maximum response (E_max_ ± SEM) of 340 ± 19 % above pre-drug levels. Its potency (pEC_50_±SEM) was 8.0 ± 0.1. The PUFA-derived eCBs DHEA, γ-LEA, AEA, and NADA also triggered TRPV1 activation. In contrast, eicosapentaenoyl ethanolamide (EPEA), 2-AG, 2-AG ether, and 2-LG showed minimal effectiveness (Table 2; Figure 1). Moreover, the response of the endocannabinoid-related metabolites EPEA, 2-AG, 2-AG ether, and 2-LG at 30 µM is significantly lower than the maximal response of capsaicin (10 µM), while the maximal responses of the others did not differ from those produced by the highest concentration of capsaicin (*Table S1*).

**Table 2:**
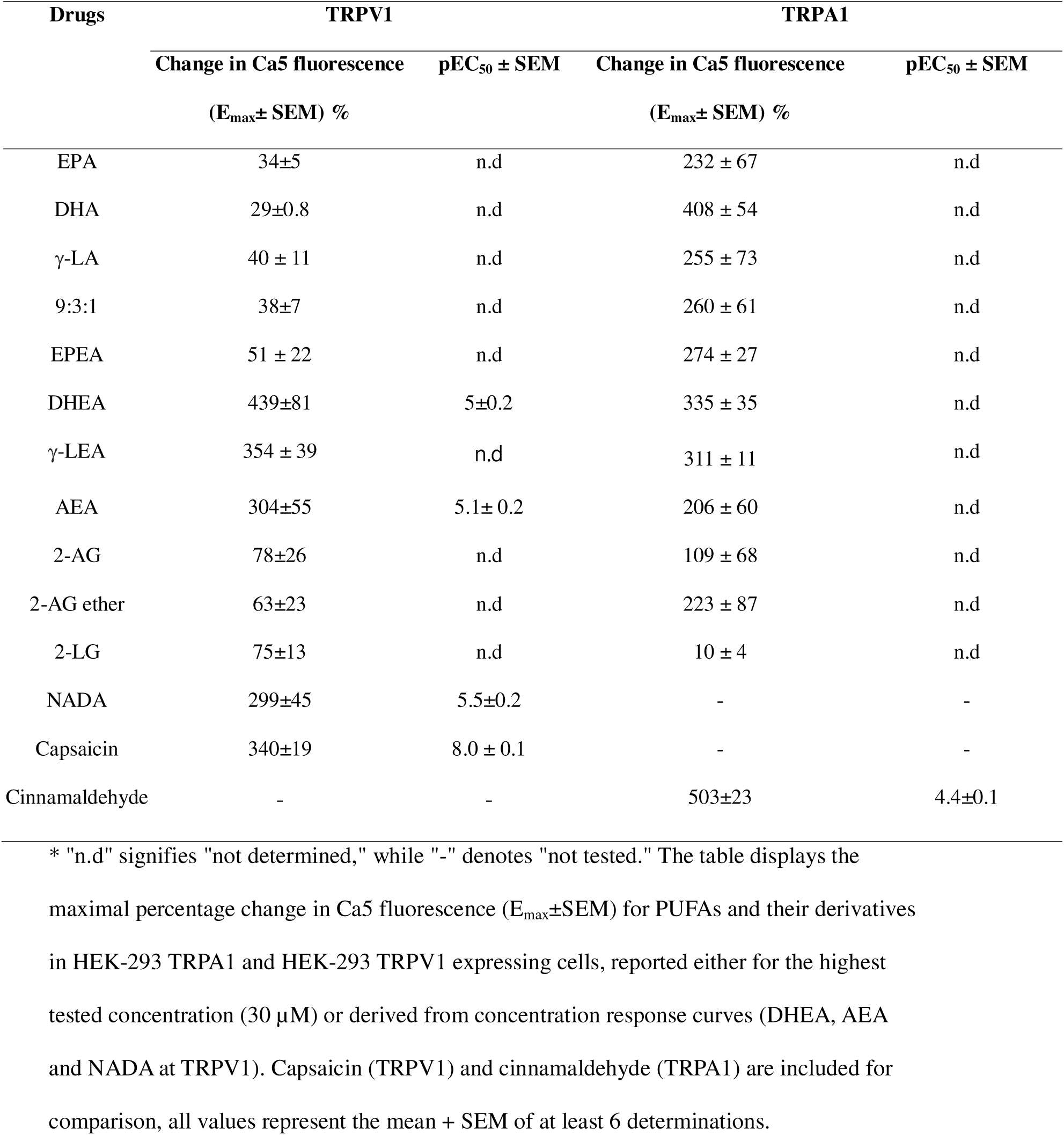
The response of endocannabinoid-related metabolites and their parent compounds in HEK-293 TRPV1 and TRPA1 expressing cells.

| Drugs | TRPV1 |  | TRPA1 |  |
| --- | --- | --- | --- | --- |
|  | Change in Ca5 fluorescence | pEC<sub>50</sub> ± SEM | Change in Ca5 fluorescence | pEC<sub>50</sub> ± SEM |
|  | (E<sub>max</sub>± SEM) % |  | (E<sub>max</sub>± SEM) % |  |
| EPA | 34±5 | n.d | 232 ± 67 | n.d |
| DHA | 29±0.8 | n.d | 408 ± 54 | n.d |
| γ-LA | 40 ± 11 | n.d | 255 ± 73 | n.d |
| 9:3:1 | 38±7 | n.d | 260 ± 61 | n.d |
| EPEA | 51 ± 22 | n.d | 274 ± 27 | n.d |
| DHEA | 439±81 | 5±0.2 | 335 ± 35 | n.d |
| γ-LEA | 354 ± 39 | n.d | 311 ± 11 | n.d |
| AEA | 304±55 | 5.1± 0.2 | 206 ± 60 | n.d |
| 2-AG | 78±26 | n.d | 109 ± 68 | n.d |
| 2-AG ether | 63±23 | n.d | 223 ± 87 | n.d |
| 2-LG | 75±13 | n.d | 10 ± 4 | n.d |
| NADA | 299±45 | 5.5±0.2 | - | - |
| Capsaicin | 340±19 | 8.0 ± 0.1 | - | - |
| Cinnamaldehyde | - | - | 503±23 | 4.4±0.1 |

\* "n.d" signifies "not determined," while "-" denotes "not tested." The table displays the maximal percentage change in Ca5 fluorescence (E<sub>max</sub>±SEM) for PUFAs and their derivatives in HEK-293 TRPA1 and HEK-293 TRPV1 expressing cells, reported either for the highest tested concentration (30 μM) or derived from concentration response curves (DHEA, AEA and NADA at TRPV1). Capsaicin (TRPV1) and cinnamaldehyde (TRPA1) are included for comparison, all values represent the mean + SEM of at least 6 determinations.

However, despite the above-mentioned metabolites of PUFA’s were observed to significantly elevate calcium (Ca) levels, we were unable to report the potency (EC_50_ values) for these drugs because we could not construct complete concentration-response curves (CRCs) for them, owing to the insolubility of these drugs at higher concentrations.

The fatty acids, EPA, DHA, γ-LA, or their combination in the ratio of 9:3:1 did not trigger TRPV1 response (Figure 2A). To test if they inhibited activation of TRPV1 ion channel, we applied a sub-maximally effective concentration of capsaicin (10 nM) after a 5 minutes exposure to EPA, DHA, γ-LA or their 9:3:1 ratio. Capsazepine (Bevan et al. 1992; Walpole et al. 1994) was used as a positive control for inhibition of capsaicin activation of the channel.

**Figure 2.**
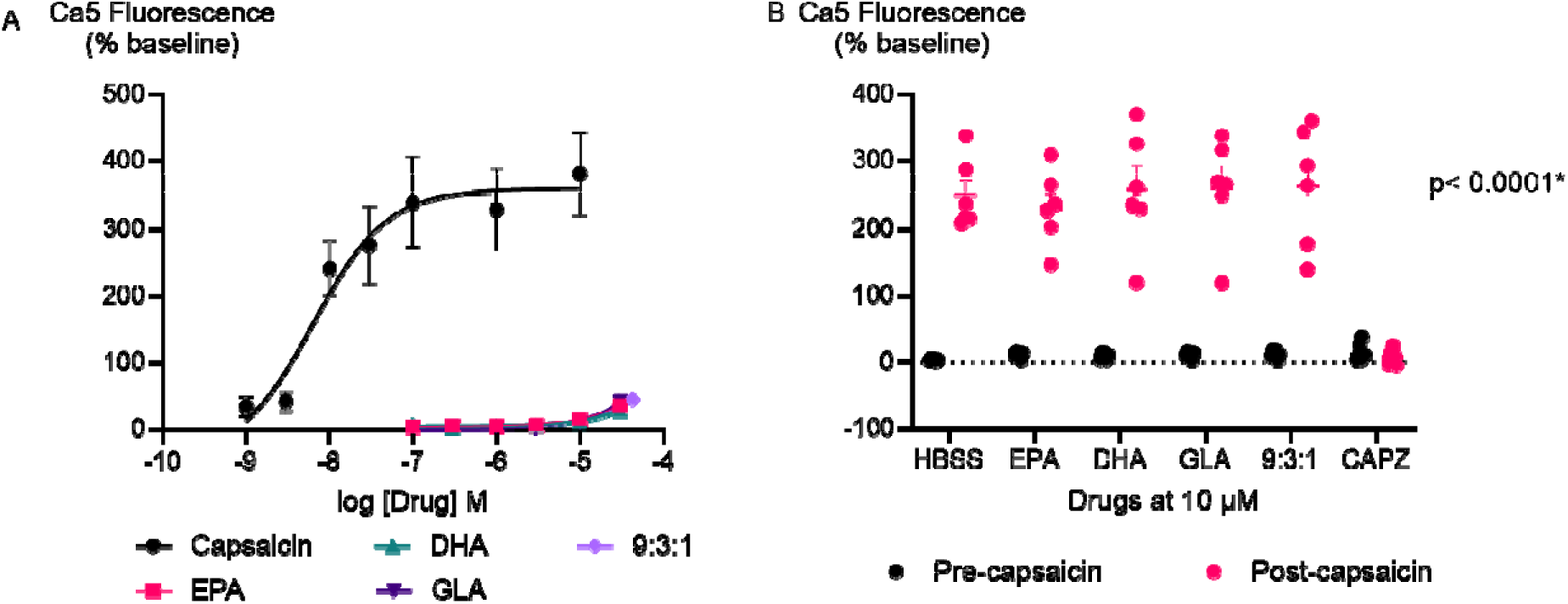
(**A)** Concentration response curves of PUFAs in HEK-293 TRPV1 expressing cells; **(B)** Effects of PUFAs on responses to capsaicin in HEK-293 TRPV1 expressing cells. PUFAs or capsazepine (10 µM each) were added to the cells for 5 minutes then capsaicin (10 nM) was added. Changes in [Ca]_i_ are expressed as percentage of the pre-drug baseline. The blue circles represent the response to first drug in individual experiments, while the red circles correspond to the response to subsequent capsaicin addition. PUFAs (10 µM) do not inhibit the subsequent response to capsaicin (10 nM), whereas preincubation of the cells with the TRPV1 antagonist capsazepine (CAPZ) (10 µM) does (P< 0.05).

Pre-incubation with capsazepine (10 µM) prevented capsaicin-induced fluorescence changes (P<0.05) (Figure 2B). However, pre-treatment with EPA, DHA, γ-LA, or their 9:3:1 ratio (10 µM) did not affect the capsaicin response in HEK-293 TRPV1 expressing cells.

Application of DHEA and γ-LEA at 10 µM as well as cinnamaldehyde at 300 µM produced an elevation [Ca]_i_ in HEK-293 TRPA1 cells significantly greater than that produced in HEK-293 EV cells. However, at 10 µM, EPA, DHA, γ-LA, their 9:3:1 ratio, as well as EPEA, AEA, 2-AG and 2-AG ether did not produce a change in [Ca]_i_ different to that seen in HEK-293 EV cells (Table 1).

The effects of the positive control and endocannabinoid-related metabolites of PUFA’s in HEK-293 TRPA1 expressing cells were concentration-dependent. Cinnamaldehyde, a commonly used activator of TRPA1, increased Ca5 fluorescence, with a maximum effect of 503±23 % above baseline with a pEC_50_ ± SEM of 4.4±0.1. EPEA, DHEA, γ-LEA, AEA, 2-AG, and 2-AG ether, also triggered the TRPA1 ion channel response, while 2-LG had no effect (Table 2; Figure 3). Among these metabolites, the response of EPEA, AEA, 2-AG, 2-AG ether, and 2-LG at 30 µM exhibited a significant difference to the response produced by the highest concentration of cinnamaldehyde (300 µM), while the maximal responses for DHEA and γ-LEA at the same concentration did not differ from it (*Table S1*).

**Figure 3.**
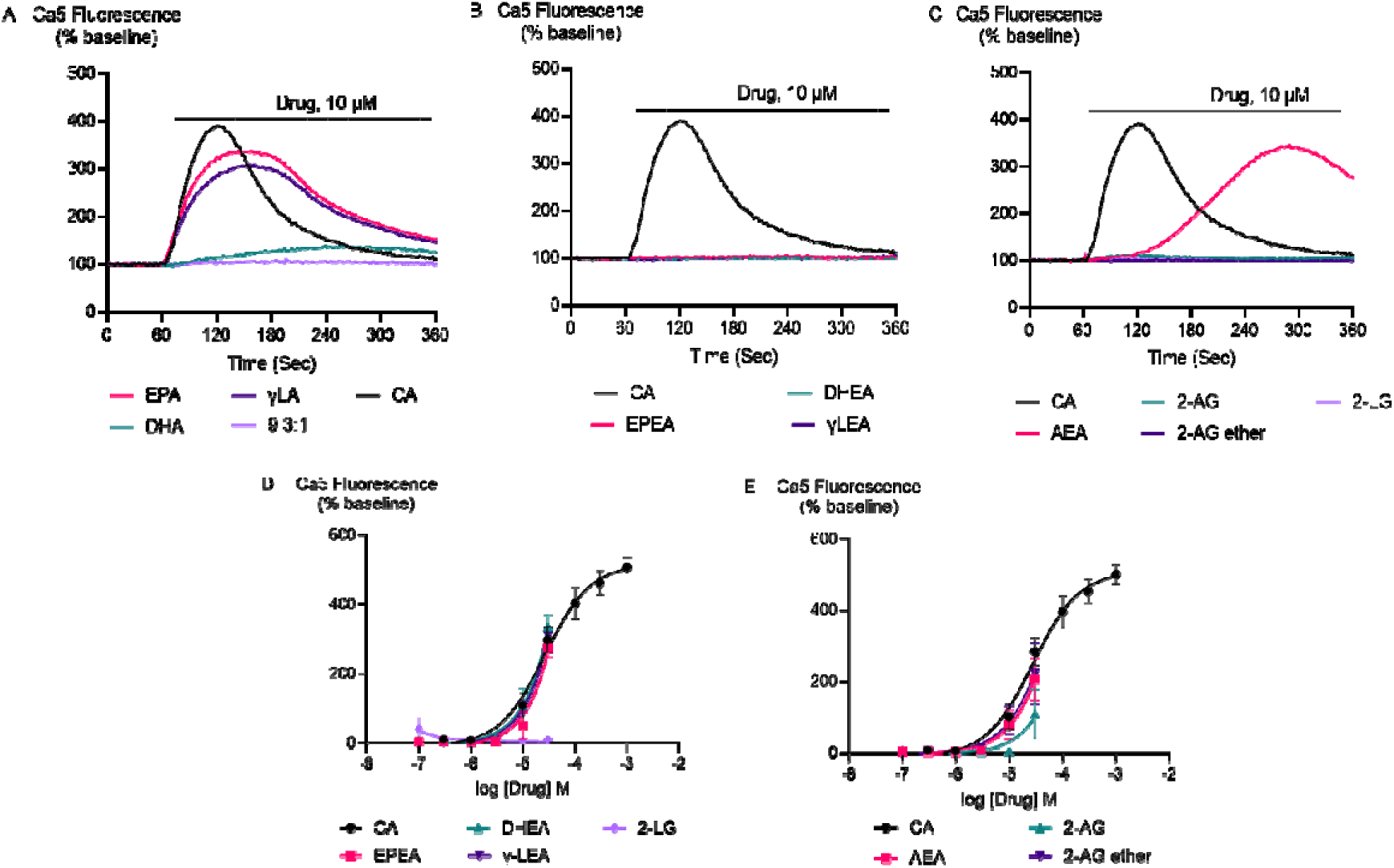
Traces of PUFAs and their endocannabinoid related metabolites at 10 µM (A, B and C) and concentration-response curves (D and E) for endocannabinoid related metabolites in HEK-293 TRPA1 expressing cells. In A), B) and C) drugs were added for the duration of the bar. The CRCs were analysed using a three-parameter logistic equation from 6 independent experiments conducted in duplicate. CA represents Cinnamaldehyde.

Furthermore, EPA, DHA, γ-LA, and their combination in a 9:3:1 ratio were found to activate TRPA1 ion channel. At the highest concentration tested (30 µM), their maximum effects were 232±67 %, 408±54 %, 255±73 % and 260±61 %, respectively (Table 2; Figure 4).

**Figure 4.**
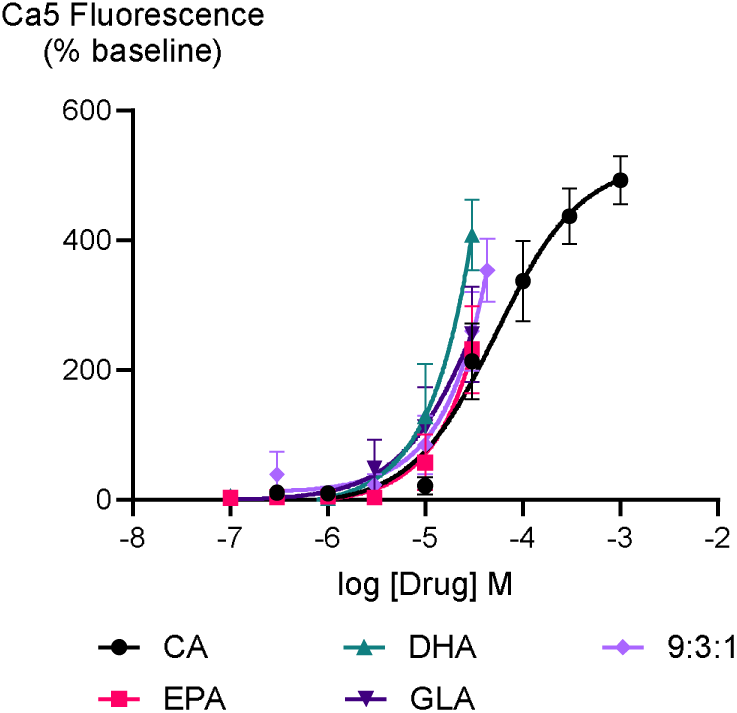
Concentration-response curves of PUFAs in HEK-293 TRPA1 expressing cells. The CRCs were fitted using a three-parameter logistic equation from 6 separate experiments conducted in duplicate. CA represents Cinnamaldehyde.

Matta et al. (2007) reported that some PUFAs, specifically EPA and DHA, activated the rat TRPV1 ion channel only following incubation with the phorbol ester protein kinase C activator, Phorbol 12,13-dibutyrate (PDBu). Therefore, we assessed whether phorbol ester-dependent stimulation of human TRPV1 response by inactive PUFA could also be observed. We initially determined the effect of phorbol 12-myristate 13 acetate (PMA), another phorbol ester that is a protein kinase C activator (Jiang & Fleet 2012), on HEK-293 TRPV1 expressing cells. Three concentrations of PMA were tested for 5 minutes followed by challenge with capsaicin at 10 nM (Figure 5A). 10 nM PMA had no effect by itself or on subsequent capsaicin responses, but 100 nM and 300 nM PMA increased [Ca]i by themselves and potentiated the subsequent response to capsaicin. However, to minimize potentially confounding effects, a concentration of 100 nM PMA was chosen. For these experiments, the response to PMA after 5 minutes was subtracted from the subsequent response to capsaicin or PUFA. PMA (100 nM) increased the effects of low, but not high, concentrations of capsaicin (Figure 5C), presumably because of saturation of the TRPV1 channels at high agonist concentrations. The response of the tested PUFAs were also increased after exposure to PMA (P<0.05 for each) (Figure 5D). To rule out the possibility that the Ca5 dye was saturated at the highest concentration of capsaicin tested, we conducted experiments with ionomycin, a calcium ionophore (Kao et al. 2010) (*Figure S2*). Ionomycin (5 µM) produced a change in Ca5 fluorescence of 673 ± 25 %, significantly greater than that produced by a maximal effective concentration of capsaicin (435 ± 15 %).

**Figure 5.**
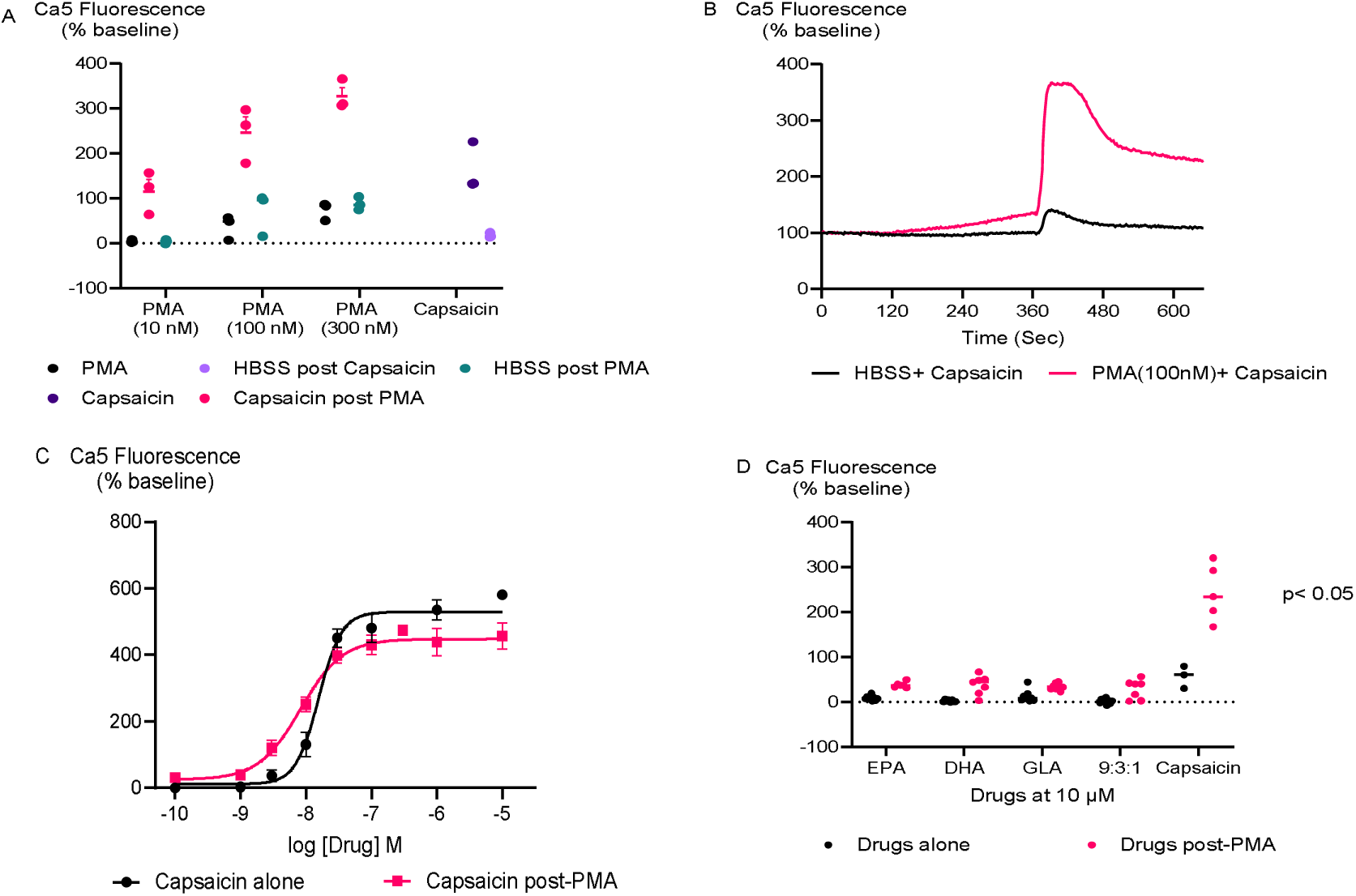
**(A)** Concentration response of PMA’s in HEK-293 TRPV1 expressing cells. PMA were added to the cells at different concentrations for 5 minutes then capsaicin was added at 10 nM. **(B)** Traces of capsaicin and **(C)** Concentration response curves of capsaicin with and without PMA in HEK-293 TRPV1 expressing cells. **(D)** Effects of PUFAs on responses to PMA potentiation in HEK-293 TRPV1 expressing cells. The cells were pretreated with PMA (100 nM) for 5 minutes then each PUFA (10 µM) and capsaicin (10 nM) was added. Changes in [Ca]_i_ are expressed as percentage of the pre-drug baseline, with the response to PMA alone at 5 minutes subtracted. The black circles represent the response of PUFAs without PMA pretreatment while the pink circles represent the response of PUFAs after PMA potentiation. The response to PUFAS was potentiated by 5 minutes pre-incubation with PMA, a protein kinase C activator (P<0.05 for each).

## DISCUSSION

The current study demonstrates that some derivatives of PUFAs directly stimulate the TRPV1 and TRPA1. Moreover, EPA, DHA, γ-LA, and their 9:3:1 ratio triggered TRPA1 activity directly, while stimulation of TRPV1 was only noted after PMA treatment of HEK-293 cells.

At 10 µM concentration, PUFAs and their endocannabinoid-related metabolites showed minimal effect on calcium levels in HEK-293 EV cells. However, significant differences were observed when comparing their effect at the same concentration between EV cells and cells expressing TRPV1 or TRPA1 receptors. To some PUFAs and their metabolites, when the concentration was increased to 30 µM, there were also notable differences in the responses between TRPV1, TRPA1-expressing cells and EV cells. These differences could be attributed to the higher concentration needed to activate TRPV1 and TRPA1 channels effectively in the TRPV1 and TRPA1 expressing cells. It is also possible that the distinct responses are due to the differential expression of TRPV1 and TRPA1 or other cellular factors in these two cell types such as differences in signalling pathways, the presence of other receptors, or variations in the cell membrane properties that affect how the cells interact with or respond to the metabolites.

Transient receptor potential vanilloid subtype 1 (TRPV1), which is widely recognized as the capsaicin receptor (Caterina et al. 1997), has previously been reported to be responsive to metabolites of PUFAs such as AEA (Zygmunt et al. 1999) and NADA (Huang et al. 2002) corroborating our findings. Our research also introduces new evidence indicating that PUFA-derived endocannabinoids (eCBs) such as DHEA and γ-LEA can trigger TRPV1 activation while EPEA showed minimal effectiveness. Moreover, consistent with our findings, previous studies have shown that 2-AG is not very effective at activating these ion channels, but it is considered a physiologically relevant activator of TRPV1 channels through phospholipase C (PLC)-mediated mechanisms and potentiation (Petrosino et al. 2016; Zygmunt et al. 2013; Zygmunt et al. 1999). Additionally, extending the findings for 2-AG, we showed that 2 related molecules, 2-AG ether and 2-LG also failed to modulate TRPV1 in these conditions.

Our findings indicate that the fatty acids EPA, DHA, γ-LA, and their combination in a 9:3:1 ratio did not trigger TRPV1 activation or inhibit its activity. As seen in earlier studies with rat TRPV1, phorbol esters enhance TRPV1 sensitivity to EPA and DHA through PKC-dependent activation (Matta et al. 2007). In our experiments, this effect was confirmed as a significant response of TRPV1 to EPA and DHA was only observed when a phorbol ester was present. Additionally, PMA amplified the TRPV1 response to capsaicin, γ-LA, and a 9:3:1 mixture of EPA, DHA, and γ-LA.

Previous reports have shown that EPA, DHA, and AEA activate TRPA1 (Ciardo & Ferrer-Montiel 2017; Motter & Ahern 2012; Redmond et al. 2014) which aligns with our findings. We have extended this work to show that, PUFA-derived endocannabinoids (eCBs) such as EPEA, DHEA, γ-LEA, and 2-AG ether can stimulate TRPA1 response, whereas 2-LG was found to be non-effective.

Moreover, it has been reported that combining ω-6 and ω-3 PUFAs is important for achieving positive health outcomes (LaChance et al. 2016; Puri & Martins 2014). Our result show that the combination of PUFAs triggers the TRPA1 ion channel activity, which may suggest a potential mechanism behind some of these observed health benefits. However, it is important to note that our study does not directly compare the effects of the combined PUFAs with those of individual components, such as EPA, DHA, and GLA alone. Therefore, it remains unclear whether the combination of ω-6 and ω-3 PUFAs in the 9:3:1 ratio is effective in activating TRPA1 than the individual fatty acids at equivalent concentrations. Therefore, future studies should aim to investigate whether the combined effects are synergistic, additive, or simply reflect the sum of their individual actions.

Additionally, it is important to consider that consuming a 9:3:1 ratio of ω-6 to ω-3 PUFAs does not necessarily translate to the same ratio in the brain or other tissues. The distribution of these fatty acids in the body may vary due to differences in absorption and metabolism, and the actual ratio reaching specific tissues such as the CNS is not well understood. Thus, ongoing research is needed to clarify how these compounds distribute in the brain and other parts of the body after consumption.

Furthermore, while TRPA1 activation plays a role in various physiological processes, its effects are not universally beneficial. For instance, activation of TRPA1 in sensory neurons is associated with pain and irritation (McNamara et al. 2007), suggesting that its activation may have different implications depending on the tissue or context. Therefore, given that TRPA1 receptors are expressed in the brain and other tissues (Nilius et al. 2012), it is essential to consider where these PUFAs and their metabolites are most likely to exert their effects. The implications of TRPA1 activation in the brain or other tissues also remain unclear and warrant further investigation to fully understand the potential health impacts.

To determine the optimal ratio of ω-6 to ω-3 PUFAs or between ω-3 PUFA’s (EPA, and DHA) for the desired health benefits, our experiment did not explore and compare other possible combinations and ratios of these compound which is recommended to be completed in future studies.

## CONCLUSION

The studied fatty acids stimulate the TRPA1 ion channel, while their metabolites trigger both TRPV1 and TRPA1 ion channels activity. Thus, local activation of these channels by PUFAs and their metabolites may influence neuronal function and provide positive effects through endocannabinoid-mediated mechanisms (Palazzo et al. 2008). Our findings indicate that these dietary components could provide neuroprotective effects by modulating these ion channels. TRPV1 channel activation is beneficial for several neuronal functions, such as regulating synaptic plasticity, influencing cytoskeleton dynamics, and aiding in cell migration, neuronal survival, and the regeneration of damaged neurons. It also integrates various stimuli involved in neurogenesis and network integration (Marsch et al. 2007; Ramírez-Barrantes et al. 2016). TRPA1 activation is also associated with modulating inflammatory responses and neuropathic pain, thereby offering protection against neuronal damage and promoting overall brain health (Nassini et al. 2014). Generally, these results highlight the therapeutic potential of dietary PUFAs in influencing brain function through specific ion channel pathways, which could support neurological health and aid in preventing neurological diseases. However, while our study provides important insights into the potential effects of PUFAs on TRPV1 and TRPA1 ion channels, the mechanisms by which orally ingested PUFAs influence brain concentrations remain unclear. Therefore, future research should focus on elucidating these pathways to better understand how dietary intake of PUFAs through supplements might alter CNS levels of these critical compounds.

## ACKNOWLEDGEMENT

This work was supported by the Australian Research Council Industrial Transformation Training Centre for Facilitated Advancement of Australia’s Bioactives (Grant IC210100040) and the Research Attraction and Acceleration program funding from the office of the Chief Scientist and Engineer, investment NSW.

## List of abbreviations

Abbreviation: Explanation
2-AG: 2-arachidonoylglycerol
2-LG: 2-linoleoyl glycerol
AA: Arachidonic Acid
AEA: Anandamide
CB1: Cannabinoid Receptor 1
CB2: Cannabinoid Receptor 2
DHA: Docosahexaenoic acid
DHEA: Docosahexaenoyl ethanolamide
DMEM: Dulbecco’s Modified Eagle’s Medium
DMSO: Dimethyl sulfoxide
eCB: endocannabinoid
ECS: Endocannabinoid system
EPA: Eicosapentaenoic acid
EPEA: Eicosapentaenoyl Ethanolamide
EV: Empty Vector
FBS: Fetal Bovine Serum
HBSS: Hanks’ Balanced Salt Solution
HEK-293: Human Embryonic Kidney cell-293
NADA: N-arachidonoyldopamine
P/S: Penicillin-Streptomycin
PPARs: Peroxisome-Proliferator Activated Receptors
PAR-1: Protease activated receptor 1
PUFA: Polyunsaturated fatty acid
TRP: Transient Receptor Potential
TRPA1: TRP ankyrin
TRPV1: TRP vanilloid
γ-LA: γ-linolenic acid
γ-LEA: γ-Linolenoyl ethanolamide
ω-3: Omega-3
ω-6: Omega-6

