## supplement table and figure for "Polyunsaturated fatty acids and their endocannabinoid-related metabolites activity at human TRPV1 and TRPA1 ion channels expressed in HEK-293 cells"

**Supplementary Table 1**: Comparison of the response of PUFA’s and their endocannabinoid related metabolites with the positive control of HEK-293 TRPV1 and TRPA1 expressing cells.

| **Drug** | **TRPV1** | | **TRPA1** | |
| --- | --- | --- | --- | --- |
|  | **Mean± SEM (N)** | **P-Value** | **Mean± SEM (N)** | **P-Value** |

| EPEA | 51± 56 | **<0.0001** | 274± 60 | **0.0183** |
| --- | --- | --- | --- | --- |

| DHEA | 306± 56 | 0.9985 | 335± 60 | 0.2155 |
| --- | --- | --- | --- | --- |
| **γ-**LEA | 354± 56 | >0.9999 | 311 ± 60 | 0.0920 |
| AEA | 229± 56 | 0.3420 | 206 ± 60 | **0.0006** |
| 2-AG | 78± 56 | **0.0001** | 109 ± 60 | **<0.0001** |
| 2-AG ether | 63± 56 | **<0.0001** | 223 ± 60 | **0.0015** |
| 2-LG | 75± 56 | **0.0001** | 10± 60 | **<0.0001** |
| NADA | 245± 56 (7) | 0.4672 | - | - |
| Capsaicin | 337 ± 56 (21) | - | - | - |
| Cinnamaldehyde | - | - | 461 ± 60 (14) | - |

* One-way ANOVA with Dunnett’s multiple comparisons was performed to compare the response of endocannabinoids and related metabolites of PUFAs at 30 µM with the highest concentration of capsaicin (10 µM) and cinnamaldehyde (300 µM) on HEK-293 TRPV1 and TRPA1 expressing cells, respectively. Unless specified otherwise, the sample size (N) is 6; a significance level of P<0.05 indicates a statistically significant difference and those values which have significant difference are written in bold format.

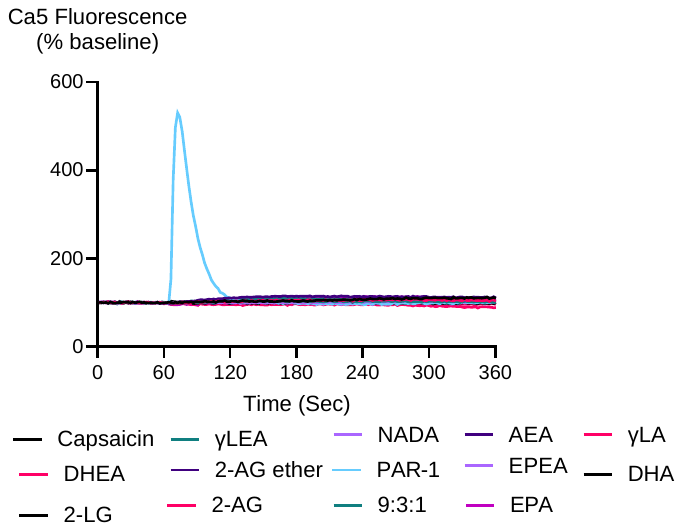

**Supplementary Figure 1.** Traces of PUFAs and their endocannabinoid related metabolites at 10 µM in HEK-293 empty vector cell.

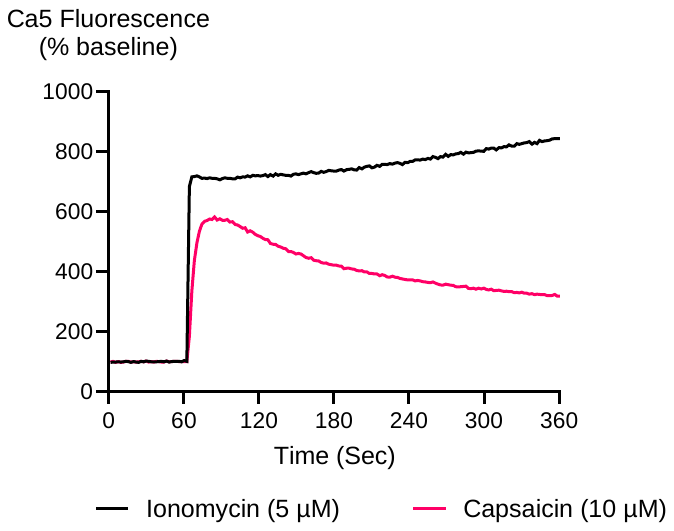

**Supplementary Figure 2.** Traces of Ionomycin and Capsaicin in HEK-293 TRPV1 expressing cells.
